# Hexagonal modulation of theta rhythmic attentional sampling of visual space

**DOI:** 10.1101/2025.02.04.636477

**Authors:** Ricardo Kienitz, Jan Martini, Randolph F. Helfrich

## Abstract

Spatial attention improves visual perception by selecting behaviorally relevant sensory signals. Traditionally, attention has been conceptualized as a static spotlight, while recent evidence posited that attention operates as a moving spotlight that samples visual space sequentially in discrete snapshots that are clocked by theta rhythms (∼3-8 Hz). While theta rhythmic attentional sampling has mainly been observed in fronto-parietal and occipital areas, theta oscillations also hallmark entorhinal-hippocampal grid-cell networks, which encode physical space in hexagonal patterns that guide overt exploration and navigation. We hypothesized that visual attention might rely on the same underlying principles and sample visual space in a hexagonal, grid-like configuration. To test this hypothesis, twenty participants performed a cue-guided attention task that probed behavioral performance as a function of space and time. Reaction times were assessed as a function of spatial location and varying cue-target intervals, which revealed prominent, spatially-structured theta rhythms. Specifically, higher theta power was evident at spatial locations that were aligned to multiples of 60°, consistent with an underlying hexagonal organization. Participants that exhibited stronger hexagonal sampling relied less on the spatial cue to guide their attentional allocation. In sum, these findings suggest that covert visual attention relies on an underlying hexagonal grid-like structure known from the entorhinal-hippocampal system and highlight that theta rhythms reflect a common organizing principle for spatial cognition.

**Significance Statement:** Attention prioritizes sensory inputs to optimize behavior. But how does attention sample the environment in space and time? Here, we demonstrate that attentional sampling of visual space is not uniform, but preferentially explores locations that are oriented along a hexagonal pattern, reminiscent of the spatial configuration of entorhinal-hippocampal grid cells. Moreover, covert attentional sampling was clocked by theta oscillations (3-8 Hz). In sum, these findings provide evidence for a shared neural basis of underlying spatial attention and navigation and reveal that theta rhythms orchestrate sampling behaviors in space and time as a unifying principle underlying spatial cognition.

## Results

Spatial attention prioritizes and selects behaviorally-relevant sensory signals to optimize visual perception that guides goal-directed behavior (Buschman & Kastner, 2015). Traditionally, spatial attention has been conceptualized as a *static* spotlight that constantly amplifies visual input at a cued location. However, more recent theories suggest that attention might operate as a *moving* spotlight that samples and explores visual space sequentially (Fiebelkorn et al., 2013; Landau & Fries, 2012; VanRullen, 2016). Critically, the sequential sampling of visual space does not occur randomly but is clocked by rhythmic brain activity. Converging evidence from behavior, (non-)invasive human and non-human primate electrophysiology studies jointly suggests that theta rhythms in the frontoparietal attention network and visual cortex orchestrate the rhythmic sampling of visual space (Fiebelkorn et al., 2018; Helfrich et al., 2018; Kienitz et al., 2018). Specifically, it had been proposed that alternating phases of theta oscillations provide distinct time windows to sample a spatial location and then shift attention to the next relevant location; hence, explaining why attention-guided visual perception is not static over time, but fluctuates as a function of the endogenous theta rhythms (Fiebelkorn & Kastner, 2019; Kienitz et al., 2022).

However, theta rhythmic behaviors are not confined to visual attention, but have been described in a variety of other sensory and cognitive modalities (Canolty & Knight, 2010; Colgin, 2013; Fries, 2023; Herweg et al., 2020; Lisman & Jensen, 2013). Theta oscillations are the most prominent electrophysiological signatures in the entorhinal-hippocampal system (Buzsáki, 2002, 2005) where their activity coordinates the firing of grid and place cells (Colgin, 2013; Moser et al., 2008). It has been firmly established that these cells represent the surrounding physical space and define a grid-like hexagonal pattern that facilitates orientation and navigation (Hafting et al., 2005; Moser et al., 2008). Similar to the theta rhythmic attentional sampling of the visual space, hippocampal theta sweeps might explore the surrounding physical space to identify the next navigational target to plan future movement trajectories (Vollan et al., 2024). Here we hypothesized that theta rhythmic attentional sampling might also sample space non-uniformly and preferentially explore locations that are oriented along an underlying spatial grid-like pattern that resembles the well-known hippocampal-entorhinal organization. Given the high degree of similarity between theta-dependent behaviors in covert and overt exploration of space, we specifically tested if attention samples the visual environment in a grid-like, hexagonally-oriented pattern.

To probe whether the rhythmic attentional sampling of visual space follows a hexagonal modulation pattern, twenty participants performed a variant of a classic spatial attention task (Buschman & Kastner, 2015; Posner et al., 1980). On every trial, participants were presented with a bar that was oriented in one of six different directions (0-150°in steps of 30°; **Figure 1A**). After a delay, a spatial cue (90% validity) indicated the likely target position. The target stimulus was presented above perceptual threshold after a variable cue-target-interval (500-1500 ms). This design enabled resolving reaction times of target detection performance as a function of space and time. We predicted a hexagonal modulation of the resulting theta rhythmic sampling behavior with stronger sampling along the preferred cardinal axis φ as well as at directions φ plus integer multiples of 60° (**Figure 1B**). In contrast, we expected lower theta rhythmic sampling along non-aligned orientations, e.g., φ + 30°.

**Figure 1.**
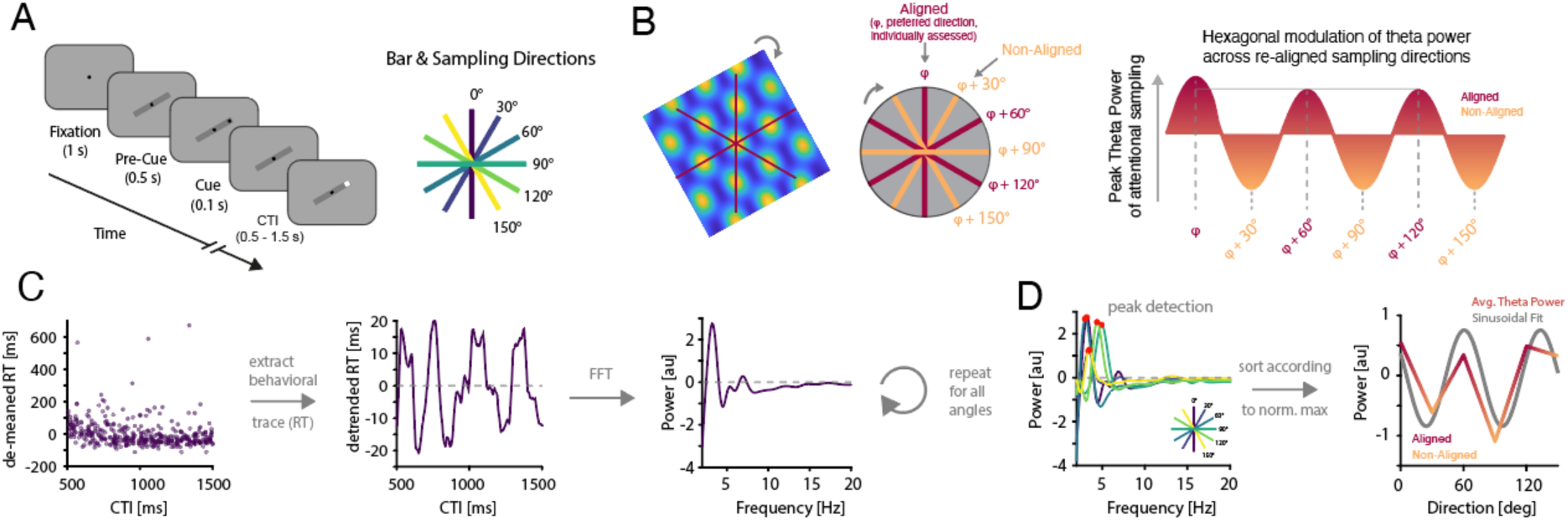
Experimental design and approach. (**A**) Left: Participants performed a spatial attention task attention task where they reacted to high-contrast target indicated an increase in luminance after having received a spatial cue (90 % validity) at either end of a bar. Different cue-target-intervals (CTI) allowed resolving reaction times as a function of time (as outlined in panel **C**). Right: The bar was randomly displayed at one of 6 possible main directions (right panel), allowing to assess attentional sampling along the 0, 30, 60, 90, 120 and 150° axes. (**B**) The experimental design was geared towards assessing a hexagonal modulation of rhythmic attentional sampling. As entorhinal grid cell organization yields stronger activation in *aligned* directions (i.e., preferred direction φ as well as φ plus multiples of 60°) compared to *non-aligned* directions, we tested if attentional sampling along different directions also exhibits such a hexagonal modulation. (**C**) Left: Single subject, single trial reaction times (demeaned). Center: Behavioral fluctuations were extracted from trials of a given orientation by binning and detrending reaction times across CTIs. Right: Time-resolved behavior was transformed to the frequency domain and power spectra were obtained per subject and bar orientation. (**D**) Left: Individual peaks in the theta range (2.5-8 Hz) were then detected for each subject and angle on the 1/f corrected power spectra. The preferred individual direction φ was defined as the angle that exhibited the maximal theta power. Right: Angles were then re-aligned with respect to φ.

As hypothesized, we observed a behavioral benefit at the cued location with significantly faster reaction times for cued than uncued targets (p = 0.03, Wilcoxon ranked sum test; 294.9 ± 8.1 ms vs. 307.9 ± 7.3 ms, mean ± SEM, **Figure S1**). Grand-average reaction times did not differ significantly across the six different bar orientations (p = 0.99, RM-ANOVA, **Figure S1**). To resolve behavior as a function of space and time, we employed a moving window approach (window size: 50ms, step size: 1ms) to obtain a time-resolved estimate of reaction times. This approach was repeated for every bar orientation separately (**Figure 1C**). Subsequently, time-resolved behavioral estimates were spectrally decomposed after applying a Fast Fourier Transform (FFT). To assess theta power as a function of bar orientation, we detected the individual theta peak (peak in the range from 2.5-8 Hz) on the 1/f-corrected power spectra for all bar directions separately (**Figure 1D**). We observed an average theta peak frequency of 4.6 ± 0.1 Hz across all participants (mean ± SEM). The peak frequency did not differ significantly across the different bar orientations (p = 0.11, one-way ANOVA).

Our main hypothesis predicted that theta power should be modulated as a function of the precise bar orientation. Given that the preferred direction φ (i.e. direction with maximum theta power) differed across participants, we re-aligned the remaining orientations with respect to the preferred direction φ for each participant (analogous to (Doeller et al., 2010; Nau, Navarro Schröder, et al., 2018; Staudigl et al., 2018)). Relative to φ, the remaining orientations were grouped into *aligned* (φ plus multiples of 60°) and *non-aligned* directions. Note that we excluded φ from all subsequent analyses of *aligned* directions as the highest theta power defined φ and therefore, would have biased the subsequent results. We then collapsed behavioral estimates along the aligned and non-aligned orientations (**Figure 2A**). On the single subject level, a non-uniform distribution of theta peak power with higher power at φ plus multiples of 60° was evident (**Figure 2B/C**).

**Figure 2.**
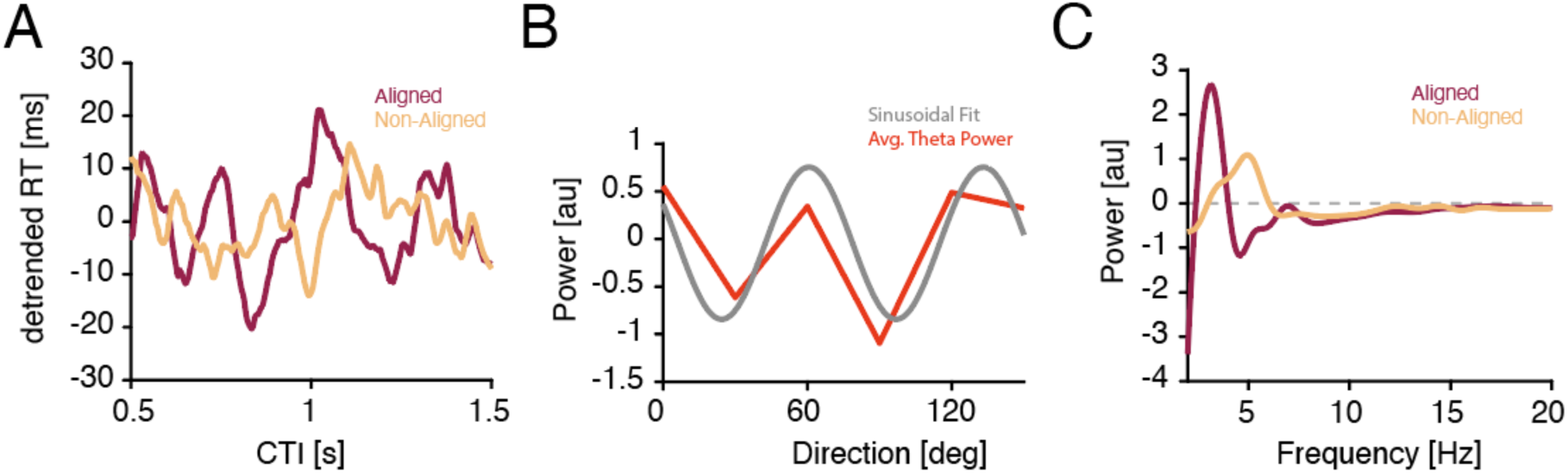
Single subject behavior. (**A**) Average detrended reaction time traces for *aligned* (red) vs. *non-aligned* (orange) angles in one exemplary participant. Note the more pronounced fluctuations of reaction times for aligned angles. (**B**) Theta peak power was non-uniformly distributed across the different angles. Higher power was found at the *aligned* (multiples of 60° with regard to φ) compared to the *unaligned* directions, indicating a hexagonal modulation across angles. (**C**) 1/f corrected power spectrum of the behavioral traces (*aligned* (red) vs. *non-aligned* (orange) directions. Note the higher power in the theta range for aligned directions. Results for this example subject implied a shift in peak frequency, which however was not present on the group level (p = 0.856, Wilcoxon signed rank test).

To quantify this observation at the group level, we repeated this analysis and alignment procedure for all participants. Across subjects, power values were non-uniformly distributed across angles (p = 9.54×10^-7^, sign-test across KL-divergences relative to uniform distributions). In the frequency domain, a clear theta peak for both, the *aligned* and *non-*aligned conditions was observed (**Figure 3A**). Yet, theta peak power significantly different between both conditions (p = 0.030, Wilcoxon signed rank test; **Figure 3B**), while peak frequency did not differ between both conditions (p = 0.856, Wilcoxon signed rank test). Critically, the theta power modulation was statistically significantly different from zero for the aligned (p = 0.021), but not for the non-aligned condition (p = 0.057). These results demonstrated that theta rhythmic sampling explored visual space non-uniformly with a preference for hexagonally oriented spatial locations.

**Figure 3.**
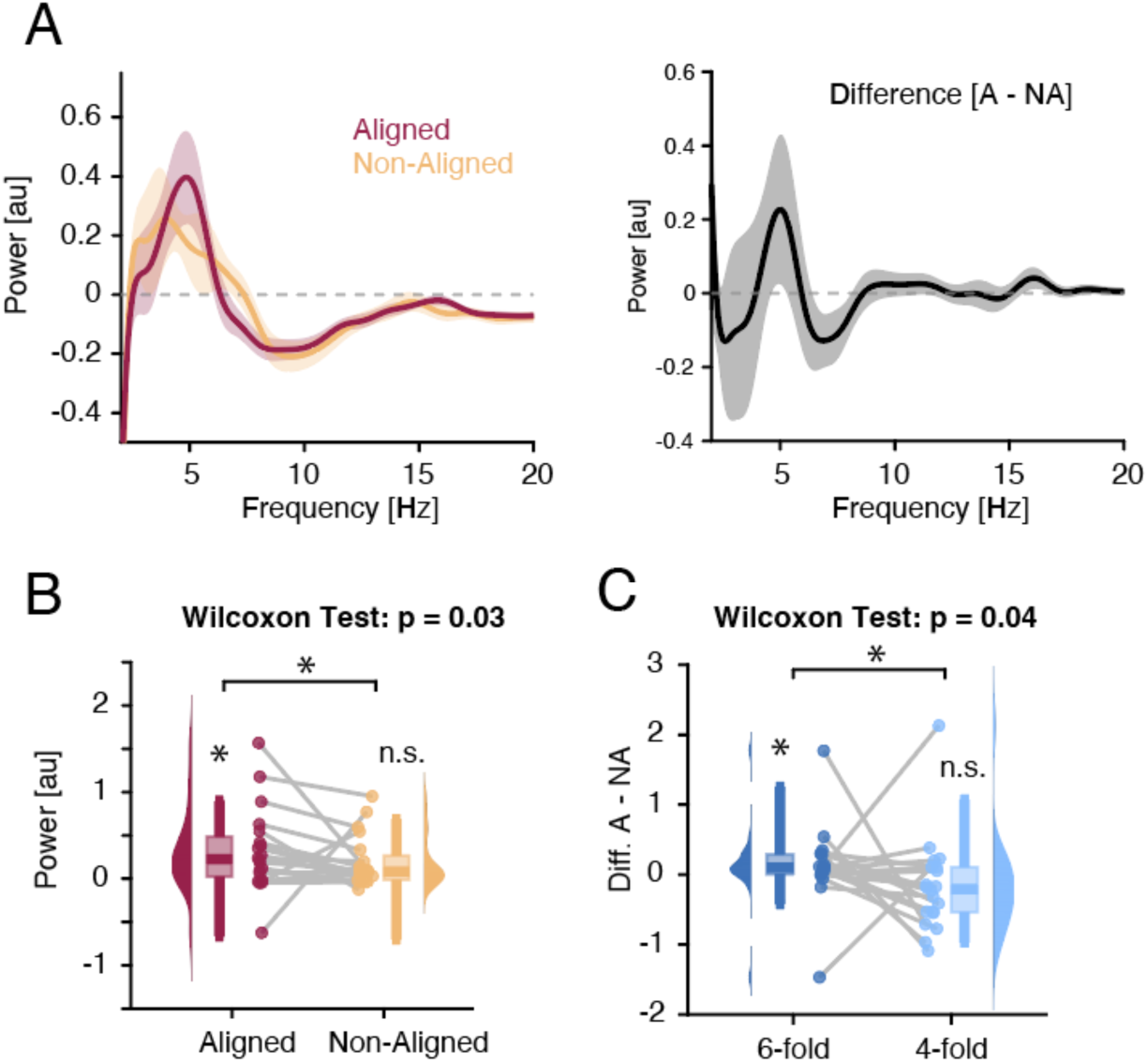
Hexagonal modulation of attentional sampling. (**A**) Left: Grand-average 1/f-corrected power spectra for *aligned* (red) vs. *non-aligned* (orange) angles. Right: Power differences between aligned and non-aligned angles across participants. Note the peak in the theta range indicating higher theta power for aligned directions. (**B**) Distributions of average theta power (4-6 Hz) for every participant for aligned (red, p = 0.021, Wilcoxon signed rank test) and non-aligned (orange, p = 0.057, Wilcoxon signed rank test) angles. Note the significantly higher average theta power for aligned angles (p = 0.030, Wilcoxon signed rank test). (**C**) Distributions of modulation strength (*aligned* – *non-aligned* angles) for every participant for a 6-fold vs. 4-fold modulation. A significantly larger modulation was observed for the 6-fold modulation (p = 0.043, Wilcoxon signed rank test). Note that only the 6-fold modulation was significantly larger than zero (p(6fold>0) = 0.008; p(4fold>0) = 0.977).

As a control, we further tested if the observed effects were specific to a hexagonal (6-fold) modulation. Hence, we repeated the analysis on surrogate data that assumed a 4-fold modulation (**Figure 3C**; *Methods*). We observed a significantly stronger modulation in the 6-fold than 4-fold scenario (p = 0.043, Wilcoxon signed rank test). Critically, a significant modulation (as compared to 0) was observed for a 6-fold configuration (p = 0.008), but not for the 4-fold configuration (p = 0.977).

Finally, we explored whether hexagonal modulation mediates a behavioral advantage during allocation of spatial attention. Given that the hexagonal modulation provides a spatial framework for integrating information across visual space, we tested how individual hexagonal modulation of theta rhythmic sampling relates to attentional benefits of the spatial cue. Hence, we quantified the attention index as an established measure of attentional allocation (Fries et al., 2001) and assessed its relationship with the hexagonal modulation (difference in theta power between aligned and non-aligned orientations, cf. **Figure 3A**). Robust linear regression revealed a significant *negative* association between the attention index and hexagonal modulation (**Figure 4**, β = - 7.76, p = 0.0379, R^2^ = 0.24; linear correlation r = −0.49), indicating that stronger hexagonal modulation is linked to a less pronounced benefit of spatial cue.

**Figure 4.**
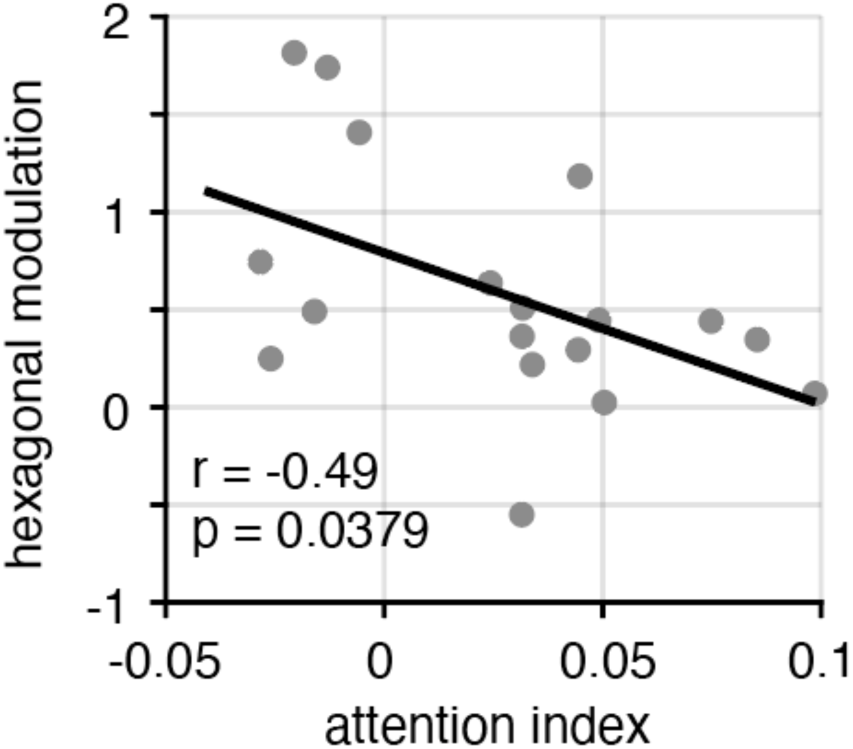
Hexagonal modulation and attentional spatial cueing benefit. Scatter plot illustrating the significant negative correlation (r = −0.49) between the attention index (higher values indicate stronger benefits of the spatial attention cue) and hexagonal sampling (higher values indicate stronger hexagonal modulation of sampling).

Collectively, these results suggest that attention-guided visual target detection preferentially operates along an underlying hexagonal configuration that is oriented in 60° steps, in accordance with a grid-like layout. Interestingly, rather than enhancing attentional benefits, stronger hexagonal modulation was associated with a reduced cueing effect; suggesting that individuals showing strong hexagonal sampling may sample visual space more extensively, and hence, rely less on a spatial attention cue.

## Discussion

Here we demonstrate that *covert* attentional sampling of visual space is not uniformly organized, but follows an underlying hexagonal structure, which mimics the structure that defines the *overt* exploration of physical space known from entorhinal-hippocampal spatial coding. Analogous to the theta-dependent exploration of the environment, we observed theta rhythmic attentional sampling of visual space that was differentially modulated across different orientations adhering to a 6-fold hexagonal structure. Hence, these results demonstrate how attention samples the visual environment in space and time and provide a perspective how covert and overt behaviors might be linked through theta rhythmic interactions.

### Theta rhythms and attentional sampling

Theta oscillations have been widely implicated in rhythmic attentional sampling, facilitating the sequential covert exploration of visual space (Fiebelkorn et al., 2013; Landau & Fries, 2012). This phenomenon spans multiple scales, from localized oscillatory activity (Kienitz et al., 2018) to large-scale network interactions in non-human primates and humans (Fiebelkorn et al., 2018; Helfrich et al., 2018). Consistent with this body of evidence, our results revealed robust theta rhythmicity in behavioral traces, evident as clear spectral peaks in the theta range in 1/f-corrected power spectra.

While previous studies established the temporal structure of rhythmic attentional sampling (Fiebelkorn & Kastner, 2019; Fries, 2023), we observed that this theta rhythmic modulation was not uniformly distributed across visual space. In the case of overt spatial exploration, it had been shown that entorhinal theta oscillations are a dominant signature and encode space in hexagonal pattern (Maidenbaum et al., 2018). These findings were recently extended by reports that overt exploration of *visual* space also show hexagonal modulations in humans (Staudigl et al., 2018). Hence, we predicted that theta oscillations might subserve a domain-general organizing principle of spatial and temporal cognition. In line with this prediction, our results revealed a non-uniform, hexagonal modulation of attentional sampling. This modulation of attentional sampling — a covert and cue-guided version of spatial exploration — is reminiscent of mechanisms that govern *overt* spatial exploration.

While spatial exploration has been studied in primates (Jutras et al., 2013; Killian et al., 2012) and humans (Doeller et al., 2010; Nau, Navarro Schröder, et al., 2018; Staudigl et al., 2018), much of the foundational work has been conducted in rodents, where place and grid cells in the entorhinal-hippocampal system are known to orchestrate navigational processes that are clocked by theta rhythms (Moser et al., 2008; O’Keefe, 1976). Moreover, recent studies in non-human primates and humans have demonstrated that grid-like signals in the entorhinal cortex can be triggered by covert spatial attention, independent of physical movement (Giari et al., 2023; Wilming et al., 2018). Our findings further substantiate this line of inquiry and demonstrate that rhythmic attentional sampling – a core principle of spatial attention – is modulated by an underlying hexagonal structure. These results suggest that rhythmic attentional sampling may rely on the same organizing principles that govern spatial exploration and navigation. In addition, our findings further imply that the observed hexagonal modulation is not a mere byproduct of rhythmic attentional sampling but may play an active role in shaping behavior. However, rather than enhancing attentional cueing benefits, stronger hexagonal modulation was associated with a reduced cueing effect, suggesting that individuals with more pronounced *intrinsic* hexagonal sampling may rely less on *extrinsic* spatial cues. This is in line with previous reports in non-human primates that demonstrated reduced neural theta oscillations in the visual cortex during focused attention (Spyropoulos et al., 2018). This raises the possibility that hexagonal modulation reflects an intrinsic spatial sampling mechanism that complements top-down attentional control.

### A common neural basis for covert and overt behavior?

It had long been speculated that attentional sampling relies on the same circuitry as overt behaviors as exemplified by the premotor theory (PMT) of attention (Rizzolatti et al., 1987). In line with a frontal origin, theta-dependent covert sampling has been observed in the frontal eye fields (FEF) as well as in adjacent frontal areas (Fiebelkorn et al., 2018; Helfrich et al., 2018; Raposo et al., 2023). Moreover, Gaillard et al. reported that saccadic eye movements are paced by a theta rhythm in FEF and explore space rhythmically (Gaillard et al., 2020). Given the prevalence of theta oscillations in fronto-parietal, occipital as well as entorhinal-hippocampal networks during both covert sampling and overt spatial exploration, it is conceivable that their characteristics rely on shared neural mechanisms. To date, it remains unresolved whether the same mechanisms given rise to theta activity in archi- and neocortex. However, there is evidence that frontal and other regions’ activity phase-lock to hippocampal theta rhythms during cognitive engagement (Hyman et al., 2005; Knudsen & Wallis, 2020; Sirota et al., 2008), thus, underscoring the notion that both are related.

Our results now provide additional behavioral evidence for a common neural basis. While theta rhythms likely originate from anatomically-distinct regions — such as the prefrontal and parietal attention network, occipital sensory areas and the entorhinal-hippocampal circuitry for navigation — they appear relevant for exploratory behaviors. Moreover, theta rhythms in different regions share several common features, such as phase coding (Kunz et al., 2019; Qasim et al., 2021; Smith et al., 2019), theta-gamma cross-frequency coupling (Canolty et al., 2006; Helfrich et al., 2018; Kienitz et al., 2021; Tort et al., 2009; Weber et al., 2024) or frequency modulation (Axmacher et al., 2010; Johnson et al., 2022) and are often reciprocally coupled (Daume et al., 2024; Johnson et al., 2023; Tamura et al., 2017). Hence, it is conceivable that theta-coupled behaviors constitute a unifying framework for spatial cognition. This consideration entails that the geometric hexagonal organization might not only subserve spatial maps, but could potentially also structure cognitive maps (Constantinescu et al., 2016; Epstein et al., 2017; Nau, Julian, et al., 2018), thus, reflecting a core principle underlying human cognition.

### Limitations and Future Directions

In the present study, we observed clear theta rhythmic sampling behavior. Critically, theta rhythmicity was modulated as a function of space. While our study provides behavioral evidence for hexagonal modulation of covert attentional sampling, several limitations need be acknowledged. First, only behavioral data was acquired, thus, we cannot resolve the putative neuroanatomical origins of the observed rhythms. It is likely that an interplay between frontoparietal attention networks, visual areas and hippocampal-prefrontal networks forms the basis for theta rhythmic sampling (Fiebelkorn & Kastner, 2019; Kienitz et al., 2022). However, previous work also implicated the thalamus in orchestrating theta-dependent network (Fiebelkorn et al., 2019; Griffiths et al., 2022; Sweeney-Reed et al., 2015). Hence, future studies that employ simultaneous, high spatiotemporal recordings from the different network nodes, ideally by means of intracranial electroencephalography (Parvizi & Kastner, 2018), need to determine whether covert and overt sampling behaviors indeed rely on the same underlying neural mechanisms.

Secondly, theta rhythmicity has been contested recently. Especially potentially inappropriate surrogate testing has been identified as a source of potential bias that disrupts inherent signal autocorrelations (Brookshire, 2022; Fiebelkorn, 2022; Re et al., 2022; Tosato et al., 2022). Here, we did not employ time-shuffled surrogate testing, but instead employed within-subject comparisons across different sampling orientations. We observed that spectral differences between aligned and non-aligned conditions peaked in the theta range (∼5 Hz). This is in line with recent lesion work that demonstrated that a focal disruption of the frontoparietal attention network alters theta rhythmic attentional sampling (Raposo et al., 2023).

Third, the employed paradigm was designed to test a specific hypothesis and thus, focused specifically on a 6-fold modulation and testing it against a 4-fold modulation pattern. Although the results support a hexagonal organization, other spatial configurations cannot be entirely ruled out. While we recorded a high trial count per participants (3000 trials across two sessions, which exceeds typically reported trial numbers, cf. (Fiebelkorn et al., 2013; Helfrich et al., 2018; Landau & Fries, 2012)) to detect subtle rhythmic effects, incorporating additional trials and spatial orientations proved challenging, since it rendered the experiment unacceptably long (> 3-4h).

Lastly, we did not include the optimal direction into our ‘aligned’ condition. Furthermore, we equated the number of angles for the ‘aligned’ and ‘non-aligned’ conditions to mitigate any sampling bias. While this is considered best practice, it needs to be stressed that omitting the optimal phase also attenuates the overall effect size.

## Conclusion

Collectively, our findings demonstrate that visual attention samples visual space along an underlying hexagonal grid-like layout. This non-uniform sampling of visual space was clocked by theta rhythmic activity. Hence, these results provide a perspective how the brain might employ similar mechanisms to support both, covert and overt exploratory behaviors, and extends known spatio-temporal coding mechanisms during spatial exploration to covert attentional processing. In sum, this study demonstrates how the brain samples the visual environment in space and time, with theta oscillations reflecting a unifying principle underlying spatial cognition.

## Materials and Methods

### Participants

20 adults (26.15 ± 4.07 years; mean ± SD; 10 females) participated in the study. The study and analyses were approved by the IRB board at the University Medical Center Tübingen (protocol number 049/2020BO2) in accordance with the Declaration of Helsinki. All participants provided informed written consent to participate in the study.

### Experimental design and procedures

Each trial began with the presentation of a central fixation point (0.7° visual angle) displayed for 500 ms to maintain participants’ visual attention. Following the fixation, a central bar, subtending 5° of visual angle, appeared for another 500 ms. The orientation of the bar was randomly chosen from 12 primary directions, equally spaced around a circle (e.g., 0°, 30°, 60°, …, 330°). After the bar presentation, a brief peripheral spatial cue, 0.7° in visual angle, was displayed for 100 ms to indicate the location where the target was most likely to appear (cue-validity of 90%). After the cue, a variable cue-target interval (500–1500 ms; CTI) was introduced. This interval was divided into 25 equal bins, with the target appearing randomly in one of these bins within each trial. The target, which subtended 0.7° of visual angle, was presented as a brief flash lasting only 17 ms. Participants were instructed to respond as quickly as possible to the target by pressing a designated key, with a response deadline of 1 second after target presentation. Responses made prematurely during the cue-target interval were recorded but marked as invalid. On 5% of trials (catch trials), no target was presented to ensure participants remained attentive to the task.

The experiment was conducted over two sessions on separate days, with each session comprising 1,500 trials, divided into 15 blocks of 100 trials each. Each of the 12 bar orientations was presented equally across trials, ensuring balanced sampling. Stimuli were generated using MATLAB (The MathWorks, Natick, MA) and the Psychophysics Toolbox and were presented on a calibrated display at an approximate viewing distance of 70 cm. Visual angles were computed based on screen dimensions and viewing distance.

### Cue Validity Effect on Reaction Times

To evaluate the influence of cue validity on reaction times, behavioral data were preprocessed to exclude outliers identified using Cook’s distance. Reaction times for valid (cue predicted target location) and invalid (cue did not predict target location) trials were averaged across participants. A Wilcoxon signed-rank test was employed to determine statistical differences between valid and invalid conditions, testing the hypothesis that valid cues yield faster reaction times.

### Reaction Time Across Target Angles

The relationship between reaction times and target angles was assessed by grouping trials based on bar orientation angles. For each participant, outliers (determined using Cook’s distance) and trials with invalid cues were excluded. Reaction times were averaged for each angle, including the angle’s counter-angle (e.g., 0° and 180°). A repeated-measures analysis of variance (RM-ANOVA) tested whether reaction times differed significantly across target angles.

### Extraction of the behavioral reaction time trace

To investigate temporal dynamics of attentional sampling, reaction time traces were computed for each participant across experimental conditions. Outliers were identified using Cook’s distance and removed alongside trials with invalid cues. For each participant, trials were grouped based on bar orientation angles, where each angle was paired with its counter-angle (e.g., 0° and 180°). Time-resolved behavioral traces were derived using a 50 ms sliding window moving in 1 ms steps, smoothed with a 25 ms window to interpolate any remaining values that resulted from the limited temporal sampling, and were aligned to the cue-target interval. Condition-specific traces were computed for each bar orientation. Time vectors were normalized to align with the CTI range and expressed in seconds for subsequent analyses.

### Spectral analysis of reaction time traces

To explore rhythmic components in reaction time data, behavioral traces were transformed into the frequency domain. To this end, we applied the Fast Fourier Transform (FFT) on preprocessed RT traces from all participants and conditions to compute power spectra for each condition and participant. Power spectra were calculated separately for all bar orientations and aggregated across trials. To correct for the aperiodic component in the power spectra, a power-law function was fitted to the frequency distribution of each spectrum, and the resulting 1/f background removed. Following spectral decomposition, grand-averaged power spectra were computed for each condition.

To investigate theta rhythmic dynamics of attentional sampling, peak frequencies and corresponding power values were extracted from the frequency-domain reaction time data. For each orientation, local maxima in the power spectrum were identified within the theta range (2.5 – 8 Hz) to determine the peak frequency and its associated power. If no clear peak was detected, the maximum power within the theta range was selected. To account for baseline fluctuations, power spectra were detrended, ensuring that periodic rhythmic activity was isolated.

Non-uniformity of power values across angles was computed as the Kullback-Leibler (KL) divergence between each subject’s observed power distribution and a uniform distribution, with a value of zero indicating no difference. We then performed a one-tailed nonparametric sign test across subjects testing whether the observed KL-divergences were systematically greater than zero. Each participant’s preferred orientation was defined as the angle with the highest detrended theta power (φ). Power and frequency values for all other orientations were then realigned relative to φ. Mean peak frequencies were calculated for each bar orientation, and a one-way analysis of variance (ANOVA) was conducted to test whether peak frequencies differed significantly across orientations.

To examine the spatial organization of theta rhythmicity in attentional sampling, power spectra were compared between bar orientations aligned and non-aligned to φ. Aligned and non-aligned angles were grouped according to a 6-fold modulation pattern (spaced by 60°), reflecting the hypothesized hexagonal organization of attentional sampling. For each participant, power spectra were averaged across aligned and non-aligned orientations, respectively. The preferred angle φ was excluded to avoid bias and number of angles was matched to avoid sampling bias. Power differences between aligned and non-aligned orientations were calculated around the prominent peak in theta frequency range (4–6 Hz, **Figure 3A**). Control analyses examined the specificity of the observed 6-fold modulation by comparing it with a 4-fold modulation pattern (spaced by 90°). A Wilcoxon signed-rank test was used to assess statistical differences in theta power between aligned and non-aligned conditions.

To examine how hexagonal modulation relates to attentional performance, we computed the attention modulation index (Fries et al., 2001), which quantifies the behavioral benefit of valid versus invalid cues. The attention index was calculated as:

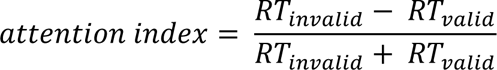

where 𝑅𝑇_𝑣𝑎𝑙𝑖𝑑_ and 𝑅𝑇_𝑖𝑛𝑣𝑎𝑙𝑖𝑑_represent the mean reaction times (RTs) for valid and invalid cues, respectively. A higher attention index reflects stronger attentional benefits, as it corresponds to a greater reduction in RTs for validly cued trials relative to invalidly cued trials. Hexagonal modulation was defined as the difference in theta power between aligned and non-aligned orientations (in a 6-fold modulation pattern) in the 4-6 Hz frequency range (cf. **Figure 3A**). We applied robust linear regression to assess the relationship between these two measures, identifying and excluding outliers using Cook’s distance (2 participants).

## Acknowledgements

This work was supported by the German Research Foundation (HE8329/2-1 to RFH), the Medical Faculty of the University of Tübingen (JRG Plus program, RFH), the Hertie Foundation (Network for Excellence in Clinical Neuroscience; RFH), the Jung Foundation for Research and Science (Ernst Jung Career Advancement Award in Medicine; RFH) and the Medical Faculty of the University of Frankfurt (Clinician Scientist Program, RK).

## Author contributions

Conceptualization: RK, RFH; Methodology: RK, JM, RFH; Formal Analysis: RK, JM; Investigation: JM; Resources: RFH; Writing – Original Draft, RK, RFH; Writing – Review & Editing: RK, JM, RFH; Supervision: RFH; Funding Acquisition: RFH.

## Supplementary Material

**Figure S1.**
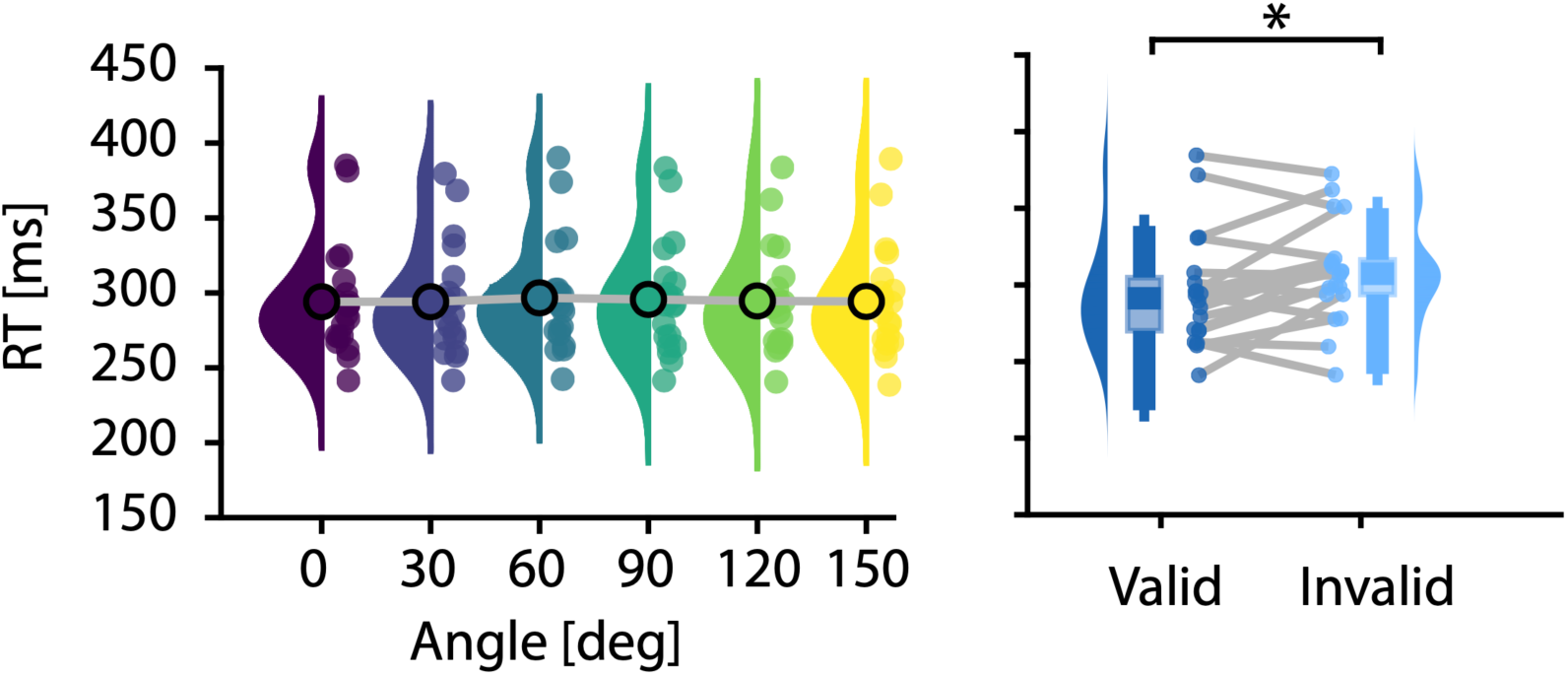
Behavioral effect of attention. Left: Grand-average reaction times (RT) across subjects did not differ significantly across the six different bar orientations (p = 0.99, RM-ANOVA). Right: Average reaction times across subjects were significantly faster for validly cued (left) compared to invalidly cued targets (right, p = 0.03, Wilcoxon ranked sum test).

